# Hypergraphs and centrality measures identifying key features in gene expression data

**DOI:** 10.1101/2022.12.18.518108

**Authors:** Samuel Barton, Zoe Broad, Daniel Ortiz-Barrientos, Diane Donovan, James Lefevre

## Abstract

Multidisciplinary approaches can significantly advance our understanding of complex systems. For instance, gene co-expression networks align prior knowledge of biological systems with studies in graph theory, emphasising pairwise gene to gene interactions. In this paper, we extend these ideas, promoting hypergraphs as an investigative tool for studying multi-way interactions in gene expression data. Additional freedoms are achieved by representing individual genes with hyperedges, and simultaneous testing each gene against many features/vertices. Further gene/hyperedge interactions can be captured and explored using the line graph representations, a techniques that also reduces the complexity of dense hypergraphs. Such an approach provides access to graph centrality measures, which in turn identify salient features within a data set, for instance dominant or hub-like hyperedges leading to key knowledge on gene expression. The validity of this approach is established through the study of gene expression data for the plant species *Senecio lautus* and results will be interpreted within this biological setting.

## 1. Introduction

In recent years multidisciplinary approaches have promoted our understanding of complex systems. In many instances, advances have been achieved by combining prior knowledge of biological systems with techniques in graph theory. The current article extends these studies by explicitly defining and applying hypergraphs for the interrogation of large data sets. We begin by rigorously establishing the background necessary to apply hypergraphs in this setting. The validity of this approach is established through the study of gene expression data, where the results are interpreted within a biological setting.

Gene co-expression networks are commonly used to study gene-to-gene interactions, and can be characterised by an incidence structure with each gene, of a genome, associated with a vertex. By studying the structure of the network, researchers seek to improve the understanding of gene regulatory networks and the underlying variation in traits of interest [1]. For instance, the network and incidence structure can be modelled with a complete graph where weighted edges record the correlation between pairs of distinct genes. That is, each vertex or gene, *g*_*i*_, is initially labelled with a vector **g**_*i*_ = [*x*_*i*1_,…, *x_in_*], recording the adjusted expression level of gene *g_i_* for *n* distinct observations. The adjustments transform the raw output of RNA sequencing, or other assays, so that comparisons can be made across genes and observations as in Section 4. Then each pair of distinct vertices (genes), ***g_i_*** ≈ [*x*_*i*1_,…, *x_ira_*], ***g_j_*** ≈ [*x*_*j*1_,…, *x_jn_*] is joined by a weighted edge, with weight,

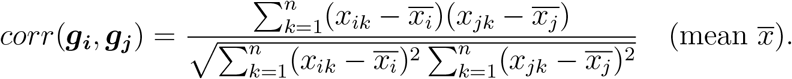

Such modelling scenarios provide valuable information about gene regulatory networks, however the following two issues exist.

First, modern methods of RNA sequencing simultaneously measure expression for a very large number of genes, resulting in dense complete graphs. As such, complete graphs may incorporate a large number of pairs of interacting genes where spurious or irrelevant correlations arise by chance, exacerbating the computational challenges. Second, weighted edges, which only capture two way gene interactions, need not reflect the known multi-way gene interactions underlying phenotypic variation in nature. While hypergraphs have been proposed to model multi-way interactions [2], such approaches may strongly compound the computational challenges [3].

We seek to move away from the gene to gene complete graph correlation analysis. Instead, we model differential gene expression using hypergraphs where vertex and edge incidence corresponds to significant expression perturbation in terms of both significance levels (*p*-value scores) and fold change for a given experimental contrast and gene. Each contrast may be a comparison between selected experimental conditions (such as a treatment and matched control), or any other parameter in a statistical model of the expression data. In this way, we may explore gene associations and multi-way interactions using hypergraphs, while reducing the complexity through the suppression of insignificant changes in gene expression levels.

The main goal of this paper is to explore and identify properties of hypergraphs, as well as a suite of visualisation instruments, that can identify and illuminate trends within complex data sets. Specifically, we will explore hypergraphs as modelling tools for the study of complex patterns within gene expression data. The benefit of this approach is that it focuses the theoretical studies of hypergraphs, motivating a refinement of hypergraph definitions and thus facilitating their application.

The definitions of applicable hypergraph properties will be provided in Section 2. It will be demonstrated that, when hypergraphs are used to model and interrogate data, basic measures such as degree centrality and cluster structure can provide valuable information about global trends within the data. Further, it will be shown that information about the intersection of hyperedges and associated centrality measures, can be used to highlight salient features of the data set. Specifically, we interrogate an experiment of differential gene expression data collected across 20 states (a factorial design with two phenotypes, two experimental conditions, and five time points) using hypergraphs. Details of these environments and the underlying experiments, together with a description of the associated data will be given in Section 3. A discussion and analysis of results will be presented in Section 4 and the biological significance of these results will be discussed in Section 5. Finally we conclude with some general observations in Section 6.

## 2. Hypergraph Analysis Tools and Definitions

This section contains relevant definitions and will use the notation 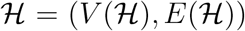 to refer to a hypergraph with vertex set 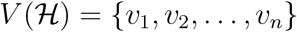 and hyperedge set 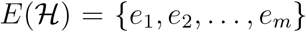, where each hyperedge in 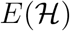 corresponds to a non-empty subset of 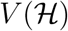, as in [4].

The number of vertices is the *order* of the hypergraph and number of hyperedges is the *size*. We only consider unweighted and undirected hypergraphs, where order is greater than zero. An example hypergraph is shown in Figure 1. The *cardinality or size* of a particular hyperedge is the number of vertices in that hyperedge, with hyperedges having multiplicity greater than or equal to one allowed. If all hyperedges have the same cardinality, say *k*, we say the hypergraph is *k-uniform*. Thus a 2-uniform hypergraph is a loop-free multigraph; if hyperedge multiplicity is at most one, it is a simple graph.

**Figure 1:**
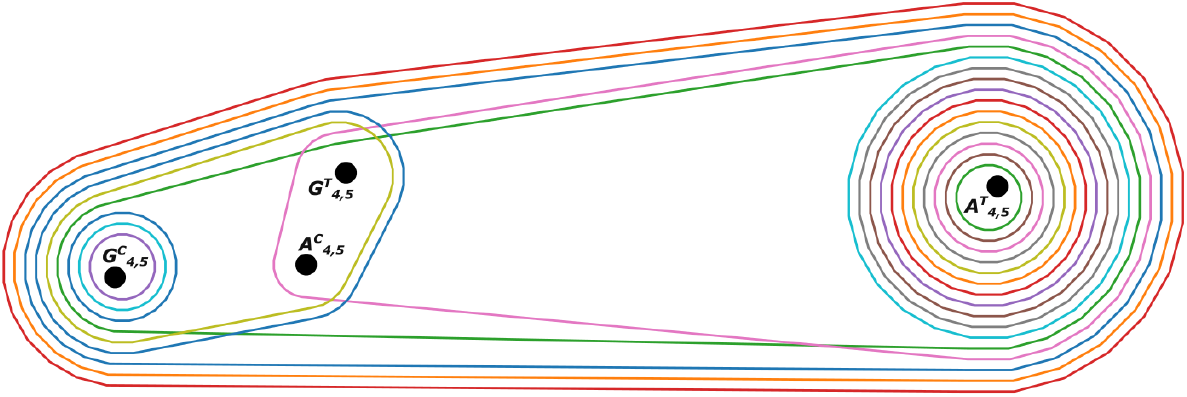
An example hypergraph of order 4, size 21 with hyperedges of cardinality 1, 3 or 4.

The hypergraph modelling approach presented is designed to interrogate a data set, consisting of a structured collection of labelled multi-dimensional data records, typically representing discrete objects. Each data record is tested against a list of conditions of interest, giving a sequence of Boolean results. The vertices of the hypergraph will correspond to the conditions and the hyperedges will correspond to the data records, with a hyperedge incident with a vertex if the discrete object satisfies the given condition. This incidence structure allows us to use the theory of hypergraphs to study the properties of the data set, see [5] for related ideas.

An incidence matrix provides a concise and unique representation for a hypergraph 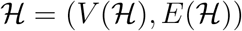, where 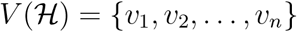 and 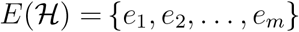. Formally the *n* × *m incidence matrix* 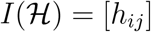 of 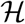 has entries defined by

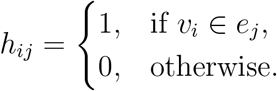

This matrix representation allows us to quickly identify the subset and number of discrete objects that satisfy a given condition and the number of conditions satisfied by a given object. Specifically, the *degree* of vertex *v*, denoted *deg*(*v*), is the number of hyperedges incident with *v*, and the *neighbourhood* of *v*, denoted *N*(*v*), is the set of hyperedges incident with vertex *v*. Two hyperedges are said to be *s-neighbours* if they share at least *s* vertices. It is clear that the sum of row *v* of 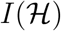 equals the degree, *deg*(*v*) = |*N*(*v*)|. For graphs and hypergraphs, the *degree distribution, P* (*κ*), which records the number of vertices of degree *κ* has been well studied [6]. If *P*(*κ*) ~ *κ*^-*α*^, where *α* > 1, we say the graph exhibits qualities of a *scale-free* graph; such graphs typically have a small number of high degree vertices, termed *hubs*, and a large number of low degree vertices, termed *spokes*. This definition also holds for hypergraphs, defining scale-free hypergraphs [7].

The degree sequence (and distribution for large order hypergraphs) can be used to study the number of discrete objects that satisfy a condition of interest and thus uniformity of the underlying data. For large and complex data sets, the order and size of the hypergraph can obscure the relationships of interest, see hypergraph 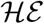 in Section 4. Thus additional methods are required to identify important structural features; we propose the use of line graphs as a primary tool.

Formally, for a positive integer *s*, the *s-line graph* 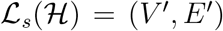 of hypergraph 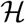, is defined by,

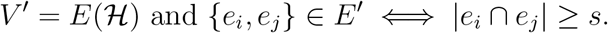

Such *s*-line graphs focus on the degree of similarity between hyperedges. The hypergraph is projected onto a simplified structure, with hyperedges represented by vertices in a simple graph. While information is necessarily lost in this construct, it allows for visual simplification and access to graph theoretical techniques, algorithms and analysis, see [8].

For clarity, hereafter we refer to vertices in the hypergraph as “conditions” and vertices in line graphs as “vertices”. The *s*-line graphs record the intersection of pairs of hyperedges and identify subsets of data points that concurrently satisfy multiple conditions. While each *s*-line graph relates to a chosen intersection size, we can explore results for various values of *s*. For a given hypergraph 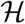, if *s* < *s*’ then the *s*’-line graph is an induced subgraph of the *s*-line graph. Typically, the *s*′-line graph would be simpler, focusing on relationships between large size hyperedges. However, this comes at the expense of excluding more of the data.

Fundamental concepts in graph theory, such as paths, distance and centrality measures can be defined using the intersection of hyperedges in a hypergraph. For instance, given a hypergraph 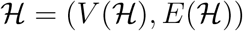 and a positive integer *s*, an *s-walk* of length *k* between hyperedges 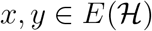 is a sequence of hyperedges in 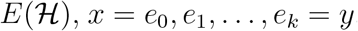, such that 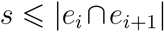 for *i* = 0,1,…,*k* – 1. This definition can be restated in terms of a list of vertices in a connected walk in the associated *s*-line graph of the hypergraph. An *s-path* then becomes analogous to a path in the associated *s*-line graph. This definition extends to *s-distance*, *d_s_*(*e,e*’), between two hyperedges *e,e*’ and is defined as the length of the shortest *s*-path over all possible *s*-walks between *e* and *e*^′^. If there is no *s*-path between two hyperedges then the *s*-distance is taken to be infinite. See [8] for more details.

If two hyperedges intersect in at least *s* conditions, then the corresponding vertices are adjacent in the *s*-line graph and thus have distance 1. On the other hand, vertices of distance 2 in the *s*-line graph identify two hyperedges that do not simultaneously satisfy *s* common conditions, however each satisfies at least *s* conditions simultaneously with a third hyperedge. This third hyperedge is said to be a *linking hyperedge*. This idea generalises with more distant vertices in the *s*-line graph indicating the distinct nature of the corresponding hyperedges.

The *diameter* of a connected graph is the greatest distance between any two vertices [9]; the diameter of an *s*-line graph is the maximum *s*-distance between a pair of hyperedges in the respective hypergraph. This provides an important summary statistic reflecting the commonality or difference between hyperedges in a hypergraph. The complete graph has diameter 1, and small but non-trivial diameters (> 1) may reflect a dense graph. In an *s*-line graph this suggests a set of at least *s* conditions that occur in most or all hyperedges. Relatively small diameters are also associated with graphs that have a hub and spoke structure, including scale free graphs. In an *s*-line graph this indicates that a minority of hyperedges satisfy many more than *s* conditions, linking pairs of hyperedges with few or no common conditions. A larger diameter *s*-line graphs implies at least one longer chain of linking hypergraphs, and the absence of either globally connected hubs or the absence of a set of *s* or more highly prevalent conditions.

Although diameter gives a foundational understanding to the structure of the hypergraph, centrality measures can provide further insight into individual hyperedges. These measures are calculated on the *s*-line graphs of a hypergraph, and hence each measure is dependent on a chosen intersection size *s*.

The first measure we consider is the closeness centrality. For a given vertex, *v*, the score is the reciprocal of the mean distance to all other vertices, so that a high score reflects shorter distances to other vertices. Formally, let *G* be a connected graph of order *n* with vertex set *V*(*G*) and *v* ∈ *V*(*G*).

Then, the *closeness centrality* score of vertex *v* is defined as,

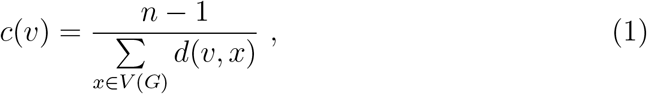

where *d*(*v,x*) defines the length of the shortest path (distance) between two vertices *v* and *x*. Since the distance between two distinct vertices is 1 or more, closeness centrality values are within the range [0,1].

By contrast we can study the centrality of a vertex by disregarding the length of these shortest paths and instead counting the number of shortest paths which traverse a vertex. In this way, the betweenness centrality of a vertex is the ratio between the number of shortest paths that are incident with the vertex and the total number of shortest paths, summed across all pairs of distinct vertices. Vertices that have large betweenness score are incident with a large number of shortest paths.

Formally, let *G* be a connected graph with vertex set *V*(*G*) and *u,v,w* ∈ *V*(*G*), then the *betweenness centrality* of vertex *v* is defined as,

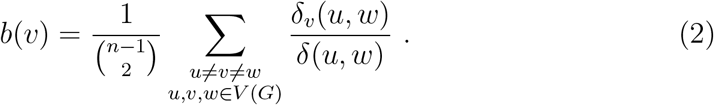

Here *δ*(*u,w*) is the number of shortest paths between *u* and *w* and *δ_v_*(*u, w*) is the number of such paths that are incident with *v*, with the first term being a normalising factor.

A betweenness score higher than the trend, can indicate a cut-vertex, a vertex that when removed disconnects the graph. In terms of line graphs, such vertices correspond to hyperedges that generate a partition of the hyperedge set. If the diameter of the hypergraph is also small, all shortest paths will be incident with a only few vertices, further biasing the measure towards cut-vertices. In this context, we can view betweenness as a measure that will aide in identifying outlier vertices and hyperedges.

The final measure that we will study is the eigencentrality. Formally, let *G* be a simple connected graph of order *n* with vertex set *V* (*G*) = {*v*_1_,*v*_2_,…, *v_n_*} and associated adjacency matrix *A*(*G*) = [*a_ij_*], where *a_ij_* = 1 if *v_i_* and *v_j_* are adjacent, and otherwise *α_ij_*· = 0. Then the Perron-Frobenius Theorem [10] states that the *dominant* (largest) eigenvalue *λ* of *A*(*G*) is positive and has multiplicity 1 and the corresponding eigenvector ***x*** is positive and unique up to scaling. Then the *eigencentrality* of vertex *v_i_* is *e*(*v_i_*) = *x_i_* [11]. By definition,

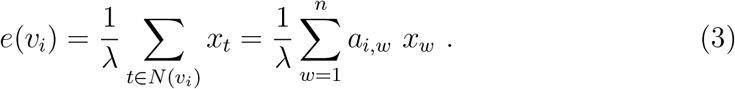

Thus this centrality measure gives a score to each vertex based on the score of its neighbours. Eigencentrality may also be interpreted using random walks on the graph. Using the set of eigenvectors {***x***_**1**_, ***x***_**2**_,…, ***x_n_***} as a basis in the expansion of the all 1 vector, *J_n_*, the power method, [11],

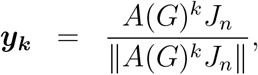

provides an iterative method for calculating an approximation for the dominant eigenvector, where for large values of *k*, ***y***_*k*_ stabilises. As such, the adjacency matrix raised to an integer index, say *k*, can be used to approximate the number of length *k* walks between vertices. In conjunction with the power method, we now associate the eigencentrality score with the frequency that a vertex occurs on a random walk of infinite length. The more accessible vertices will be traversed more often, indicating central vertex behaviour, thus being a useful tool in identifying hubs.

For the data studied here, line graphs tend to contain hubs or clusters of hyperedges, and so have small diameter (see Section 4). In this context it is worth emphasising some of the characteristics of the closeness, betweenness and eigencentrality measures with respect to small diameter graphs.

Broadly speaking, in graphs of small diameter the centrality measures will tend to increase with degree [12]. To compensate for this effect, we plot each measure against degree and avoid over-plotting by consolidating equivalent hyperedges using a colour scale to reflect multiplicity. Throughout we shall use the term *x-multiplicity hyperedge* to reflect *x* equivalent hyperedges with identical conditions. However, note that degree, size and other metrics such as centrality measures allow multiple equivalent hyperedges; or equivalently, distinct hyperedges are weighted by their multiplicity.

Further, for a graph of order *n* and diameter two, a vertex of degree *d* is distance 1 from *d* vertices and distance 2 from *n* – 1 – *d* vertices. This implies that closeness is a deterministic function of degree. Thus closeness centrality is analysed only for graphs of diameter greater than two.

It will be demonstrated that the betweenness and eigencentrality can reveal distinct information. The betweenness centrality will, for example, distinguish cut-vertices of large degree. A cut-vertex in a line graph corresponds to a hyperedge that shares disjoint subsets of conditions with separate clusters within the hypergraph and respectively the data set. Further, in diameter two graphs, a vertex of large degree and zero betweenness indicates a large cluster of hyperedges that all satisfy a set of common conditions. Since the eigencentrality of a vertex (hyperedge) is proportional to the eigencentrality of its neighbours, this measure tends to highlight subsets of vertices (hyperedges) all of high degree, emphasising subsets of strong connectivity, central within the graph and hypergraph.

For brevity in subsequent discussions, we suppress the term centrality when referring to closeness and betweenness.

## 3. Differential Gene Expression Data Set

The applicability of hypergraphs to model gene co-expression data will be explored through differential gene expression recorded over a short time series, under two distinct biological phenotypes, both tested in the presence or absence of an experimental treatment.

Specifically, we use hypergraphs to study gene expression data for the plant species *Senecio lautus*, an Australian coastal plant. This plant has adapted to diverse and changing environmental conditions, evolving certain traits and phenotypes under natural selection. In particular, in sand dune environments the ecotype consists of tall plants with an erect growth habit; while in headland environments, *S. lautus* has a low-lying growth habit, known as prostrate [13]. The trait studied in this expression experiment is *gravitropism*, where a plant will respond specifically to the pull of gravity by re-orientating its growth vertically after perturbation [14]. This trait responds strongly in the dune ecotype (*gravitropic*) but weakly, or not at all, in the headland ecotype (*agravitropic*). These responses have repeatedly evolved in independent populations of (*S. lautus*) implying that deterministic forces are responsible for their origin [13].

To understand gravitropic processes, an experiment was designed (Table 1) where 600 individual seedlings were subjected to a 90° plant rotation. Details of the data production and standard analyses of the experiments considered in this paper will be published elsewhere (Broad et al. in preparation). Here we provide an overview of the approach taken so the hypergraph analyses can be placed in a biological context. Briefly, gene expression was recorded over five time periods (30, 60, 120, 240 and 480 minutes post intervention) during which the plants may return to vertical growth patterns. Matched controls were tested in parallel, where the plants were not rotated to account for expression changes due to intrinsic differences and circadian rhythm. The measured individuals were hybrids resulting from inter-crossing Dune and Headland plants for 11 generations. This process led to the decoupling of genetic associations between traits so any signals arising from the current analyses can be ascribed to gravitropism and not other traits. Hybrid individuals were divided into groups distinguishing the 10 highest and 10 lowest gravitropic families. Hereafter we refer to these as gravitropic (*G*) and agravitropic (*A*) groups, respectively. Stem tissue from each family at each of the five time steps and for each experimental level was harvested from each gravitropic family and each agravitropic family. For each of the four experimental levels, the 10 stems were aggregated for each sampled time and sequenced with RNA-Seq, giving expression readings for candidate gene sequences. The entire experiment was carried out three times (using the same plant families), giving a total of 20 experimental states and 60 samples analysed, as summarised in Table 1.

**Table 1:** Summary of expression experiment. *N* is the total number of plants harvested of the planned 150.

| *Phenotype* | **Agravitropic (*A*)** | **Gravitropic (*G*)** |
| --- | --- | --- |
| **Control (*C*)** | $N = 144$, 10 *A* families,<br>5 times post intervention,<br>3 biological repeats | $N = 134$, 10 *G* families<br>5 times post intervention<br>3 biological repeats |
| **Rotation $90^\circ$ (*T*)** | $N = 145$, 10 *A* families,<br>5 times post intervention<br>3 biological repeats | $N = 137$, 10 *G* families,<br>5 times post intervention,<br>3 biological repeats |

The 60 pooled RNA samples were sent to Beijing Genomics Institute (BGI) Australia for library preparation and sequencing using the DNBSEQ method. The raw reads for each sample were run through FASTQC [15] to determine the quality of the reads, identifying poor-quality bases for trimming. All low-quality bases were found to be within the ten base pairs (*bp*) from the starting end, and so such regions were trimmed using the trimmomatic tool [16], resulting in reads of length 140bp. Then RNA sequencing reads were assembled and mapped onto a S. lautus reference transcriptome, with HISAT2 [17] used to map the reads onto a previously created PacBio genome [18]. The files were converted using Samtools [19] the transcripts per million (TPM) were calculated using a a TPM calculator [20]. Ultimately GMAP/GSNAP [21] was used to translate files into GFF3 format for further analysis.

A number of sequences include gene variants arising from differences between individuals, from differential splicing, and gene duplication. We collapsed sequences that mapped to a single region of the *S. lautus* into candidate sequences as genes, and we report the expression patterns for them. The number of candidate gene sequences across all categories was reduced from 894,935 overlapping sequences to 269, 210, by collapsing regions with 95% sequence overlap (intersection/union) and removing sequences with less than 10 reads in any of the 60 samples. This number of genes corresponds to an average of 13, 460 genes per experimental level. A large number of genes also had small variations in the sequence, known as alleles. The expression of these alleles was measured independently. They were not grouped to avoid losing variation, as well as to understand whether one particular allele was responding to treatment. Within the data set the alleles are numbered as “paths”. In Section 4, some of these are grouped together due to the nature of the hypergraph analysis method. Standard methods were applied to normalise expression data across the samples. Expression data is non-negative, with random variation generally proportional to observed values. The analysed data revealed heterogeneity across batches, so an average was taken for each gene at each time point across the three batches and 10 families. This is in line with standard practice in the field of functional genomics [22]. The results were collated in a file consisting of 269, 210 columns (gene variants) and 20 rows (one row for each of the time steps *t*_1_ = 30, *t*_2_ = 60, *t*_3_ = 120, *t*_4_ = 240 and *t*_5_ = 480 across each of the four groups, agravitropic control (*AC*), agravitropic treatment (*A^T^*), gravit-ropic control (*GC*) and gravitropic treatment (*G^T^*) with entries recording the relevant expression level in units of transcripts per million (TPM).

A common threshold for significant gene expression differences between treatments is generally assessed as a ratio, with 2-fold change a common threshold or benchmark for significant change [23]. Thus we applied a log_2_ transformation to the normalised expression values, then used linear models to estimate differences across time and between experiments in terms of *log*_2_-fold change. More precisely, let 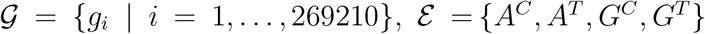, 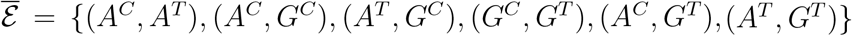, and *T* = {*t_j_* |*j* = 1,2, 3, 4, 5}. Then for gene 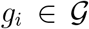, experiment 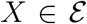 and time step *t_j_* ∈ *T*, let *g_i_*(*X,t_j_*) represent *log*_2_ of the mean normalised expression level for gene *g_i_*.

Next we defined a set of 16 conditions 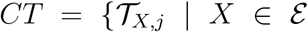, *j* ∈ {2, 3, 4, 5}}, where for 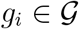,

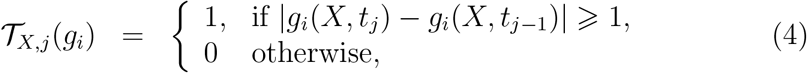

and a set of 30 conditions 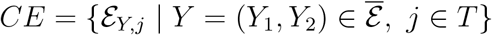, where for 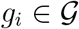,

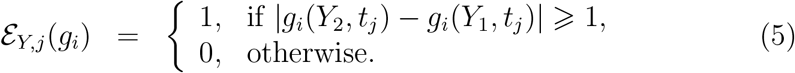

That is, the mean difference between groups or subgroups is represented by an estimated *log*_2_-fold change between them. In this way, we have a list of 46 insightful conditions, which will form the basis for the incidence structure for two distinct hypergraphs. We must be aware of the distinction between the hypergraph conditions and the experimental conditions or states in the underlying data; in standard scientific terminology, each of the 46 hypergraph conditions corresponds to a contrast between two of the 20 experimental conditions.

To control the family-wide false discovery rate in the face of large gene sets with low rates of true expression difference (the agravitropic and gravitropic groups are genetically similar), we used an empirical Bayesian approach, as implemented in the R package limma [24], to incorporate dispersion information from the full set of genes. This allowed an optimal estimate of the variance of individual results. The fitted dispersion model was used to produce a *p*-value for each condition and gene. For each condition, we then apply the Benjamini-Hochberg ([25]) procedure to the set of *p*-values across either the complete set of genes or a pre-selected set of interest, in order to produce an adjusted *p*-value for each result. For example, at most 5% of the results with an adjusted *p*-value below 0.05 should be false positives. Thus for each gene 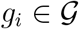, and each condition *C* ∈ *CT* ∪ *CE*, let *p*(*g_i_, C*) represent the associated *p*-value. Then we introduce an indicator function *ρ_i_* which is a mapping, 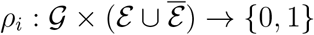, such that,

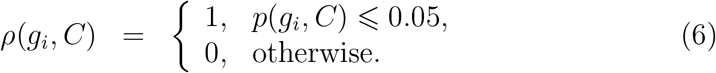

Then we define an *undirected and unweighted hypergraph* 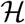, with vertex set 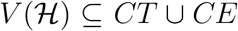 and hyperedge set 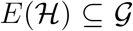, by an incidence structure where for all 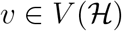 and 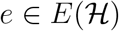 the following holds,

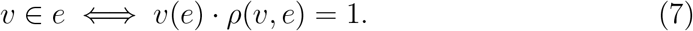

To simplify language, for the remainder of this paper we will say that a hyperedge contains a condition, if gene *g* shows significant perturbation, that is there is a 2-fold change in gene expression levels and a corresponding *p*-value less than 0.05, with respect to condition *C*.

Figure 1 provides an example of an induced hypergraph where the hyperedges correspond to a subset of 1330 gene variants, broadly classified as *transcription factor genes* and the vertices are 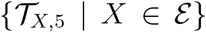 as set out in Equation (4). That is, hyperedges align with genes that have a *p*-value less than 0.05 and *log*_2_-fold change from time period *t*_4_ to *t*_5_. The hypergraph demonstrates that with respect to time steps *t*_4_ to *t*_5_, there are only four genes that exhibit significant perturbation across all four experiments, with a further two genes exhibiting significant perturbation across three of the four experiments. Further, the agravitropic treatment vertex is seen to dominate the hypergraph, with all none-empty hyperedges containing that vertex.

In the main, the examples used in this paper to study the use of hypergraphs as modelling tools will be drawn from two specific hypergraphs, 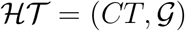 and 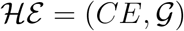.

## 4. Discussion, Analysis and Results

Our aim is to study hypergraphs as a tool for interrogating data, identifying structure and properties that highlight trends in the associated data set, exposing outliers and data points worthy of further investigation. Specifically, we illustrate how intersecting hyperedges, centrality measures, and clusters can be used to highlight salient features within a data set.

We study this in the context of the gene expression data set introduced in Section 3, with hypergraphs, as specified in Equation (7), providing the basis for our analysis. It must be emphasised that we do not present a complete analysis of this data set. We have chosen salient examples that illustrate the techniques under discussion; in this way we demonstrate the strengths of hypergraphs as interrogation tools for multidimensional data set analysis.

We will separate the set of conditions into the two types, *CT* and *CE*. The first hypergraph 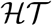, with vertex set restricted to conditions 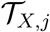 as given in Equation (4), compares expression values at consecutive times (*CT*) within each experimental group. The second hypergraph 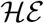, with vertex set restricted to conditions 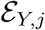 as given in Equation (5), compares expression levels across experimental groups but at the same time step (*CE*).

### 4.1. Hypergraph Summary Statistics

For large data sets where the corresponding hypergraph is of significant size (≽ 1000), visualisations as in Figure 1 are of little value. Instead, summary statistics, such as degree sequence and the hyperedge size distribution, provide an initial understanding of the hypergraph structure. These summary statistics can inform on the occurrence of (a)typical degree and hyperedge sizes, in terms of either relative constancy or else domination of key conditions or hyperedges.

The existence of a large number of conditions of low to zero degree has the obvious interpretation of little change across most data points, whereas only a few conditions of low or high degree may identify outliers. The conditions of high degree may then act as hubs, with the given conditions highlighting areas of common significant change.

**Table 2:** Degree sequence with associated information for hypergraph 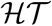.

| Condition | Group | Time 1 | Time 2 | Degree | Histogram |
| --- | --- | --- | --- | --- | --- |
| $A_{1,2}^C$ | AC | 30 | 60 | 0 | |
| $A_{2,3}^C$ | AC | 60 | 120 | 11 | █ |
| $A_{3,4}^C$ | AC | 120 | 240 | 8 | █ |
| $A_{4,5}^C$ | AC | 240 | 480 | 526 | ██████████ |
| $A_{1,2}^T$ | AT | 30 | 60 | 0 | |
| $A_{2,3}^T$ | AT | 60 | 120 | 0 | |
| $A_{3,4}^T$ | AT | 120 | 240 | 2 | █ |
| $A_{4,5}^T$ | AT | 240 | 480 | 639 | ██████████████████ |
| $G_{1,2}^C$ | GC | 30 | 60 | 0 | |
| $G_{2,3}^C$ | GC | 60 | 120 | 0 | |
| $G_{3,4}^C$ | GC | 120 | 240 | 1 | █ |
| $G_{4,5}^C$ | GC | 240 | 480 | 488 | ██████████ |
| $G_{1,2}^T$ | GT | 30 | 60 | 0 | |
| $G_{2,3}^T$ | GT | 60 | 120 | 0 | |
| $G_{3,4}^T$ | GT | 120 | 240 | 0 | |
| $G_{4,5}^T$ | GT | 240 | 480 | 390 | ██████████ |

In hypergraph 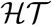, with degree sequence summarised in Table 2, the majority of conditions have degree zero, aligning with most genes showing little significant perturbation in expression level across time steps one to four, with a few exceptions. The exceptions highlight outliers where 11 or fewer genes (hyperedges) satisfy each of conditions 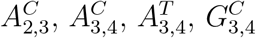. By contrast, in each of the experiments, there exists a dominant condition 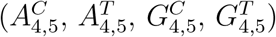 of high degree indicating that a large number of genes exhibit significant change in expression levels from time step four to five. This is perhaps not surprising given that the developmental response, gravitropism, requires time to manifest.

The hypergraph 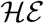 is more complex. The degree sequence (Table 3) reveals that many conditions (9 of 30) have zero degree, indicating no significant differentiation in expression levels between experiments *AC* and *AT* across all time steps. Similarly for experiments *GC* and *GT*, with the exception of time step five, where the high degree condition *G*^C^5*G^T^* indicates 371 genes exhibiting differential expression levels in this final period. The nature of this condition and these genes will be explored further in the subsequent analysis, but they suggest that the agravitropic group might be insensitive to rotation whereas the gravitropic group responds to the pull of gravity, and this is strongly visible at the end of the experiment.

**Table 3:**
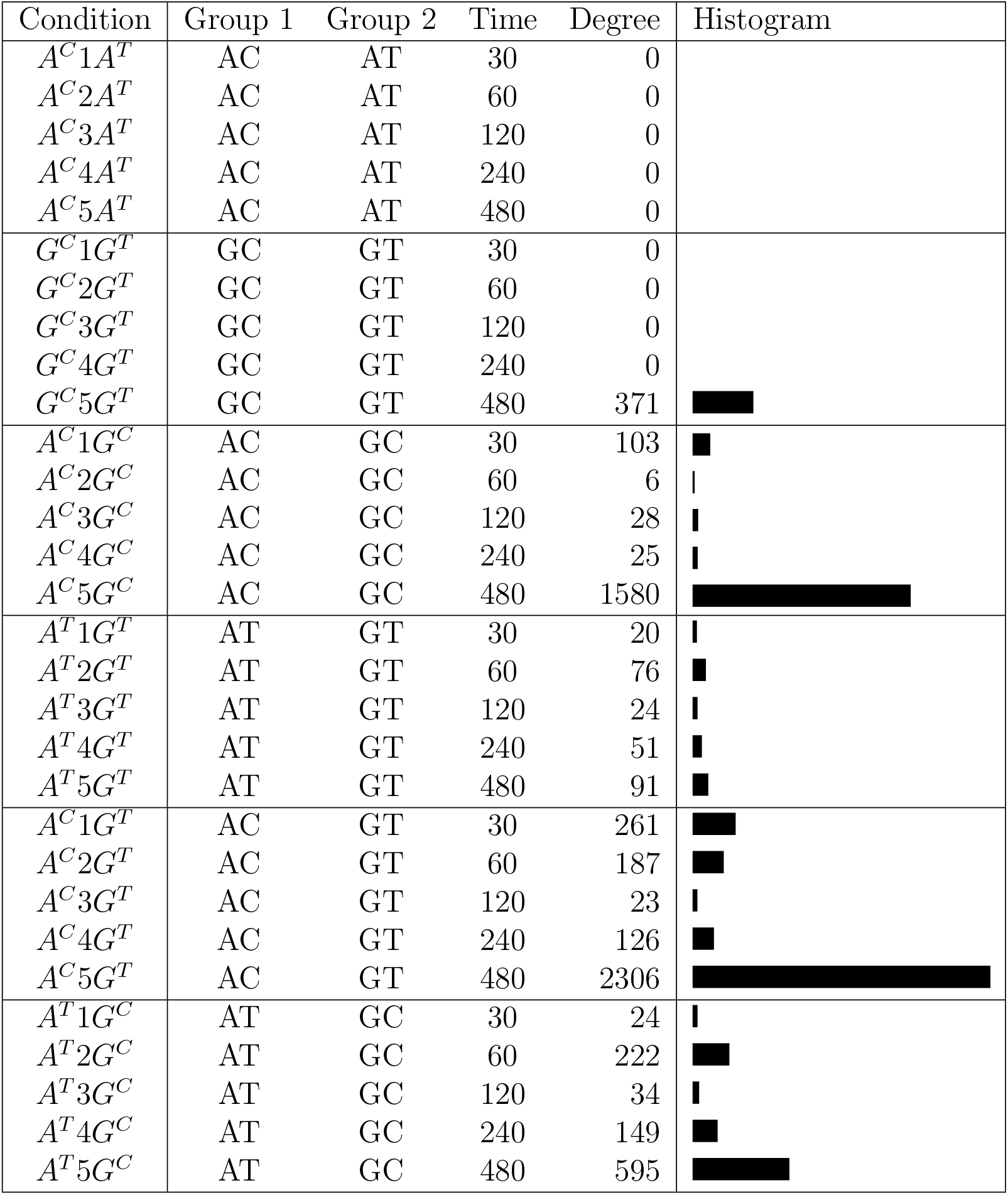
Degree sequence with associated information for hypergraph 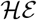.

The trend for increased differentiation in expression levels in the final time period is present across all other experimental comparisons; see Table 3 for details. The foremost delineation is for the high degree condition *A^C^*5*G^C^* incident with at least a third of the 4803 non-empty hyperedges and condition *A^C^*5*G^T^* incident with at least a half of the 4803 non-empty hyperedges. A first interpretation of these patterns is that the agravitropic control experiment is substantively differentiated from both the gravitropic control and treatment experiments, but it is not clear if these are disjoint or intersecting sets of genes (hyperedges).

Because the degree sequence revealed the conditions that show high and low contrast, but provided little information about individual conditions or patterns across groups of conditions, we investigate the distribution of hyperedge sizes to probe other salient features of the data. However, we must exercise caution, these deductions need to be viewed in light of the data and carefully interpreted.

Tables 4 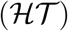 and 5 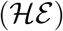 summarise the number of differentially expressed conditions satisfied by each gene through the size of the hyperedges. For both hypergraphs, the majority of hyperedges are empty, inline with expectation that only a small fraction of genes will typically exhibit differential expression across a given pair of experimental conditions. Further, the false positive rate was controlled by the multiple-testing correction procedure but at the expense of the false negative rate. That is, for a gene found to satisfy only a single condition, we cannot say with confidence that it does not have differential expression for any other contrasts. Thus the high frequency of hyperedges of size one, and the rapid reduction in counts for size greater than one, should not be over-interpreted. Nevertheless, hyperedges of size two or more do provide an improved understanding of the structure of the hypergraph and data set.

**Table 4:**
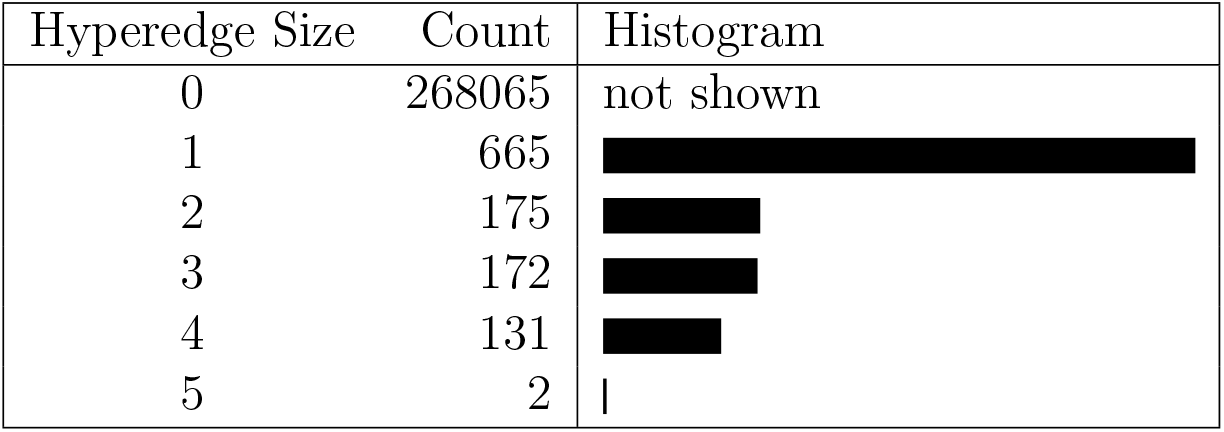
Hyperedge size distribution for Hypergraph 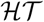.

| Hyperedge Size | Count | Histogram |
| --- | --- | --- |
| 0 | 268065 | not shown |
| 1 | 665 |  |
| 2 | 175 |  |
| 3 | 172 |  |
| 4 | 131 |  |
| 5 | 2 |  |

The relative stability in the number of hyperedges of size two, three and four in Table 4 tends to suggest that genes (hyperedges) that satisfy two conditions are likely to satisfy three or four. Combining this knowledge with the degree sequence in Table 2 suggests that genes that satisfy one of the four conditions, 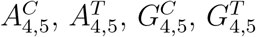 are likely to satisfy 2, 3, or all 4 of these conditions. We shall investigate this further.

Additional information available through Table 4 is the existence of two outlier genes which satisfy five conditions and Table 5 indicates the existence of a small number of hyperedges of sizes from 7 to 20 at the tail of the hyperedge size distribution. Such hyperedges are worthy of further investigation, with the potential for identifying genes of higher biological significance within this experimental setup. For instance consider the hyperedge of size 20 in 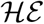. There are 21 conditions with non-zero degree, and interrogation of the data shows one condition *G^C^*5*G^T^* of non-zero degree that does not lie on the hyperedge of size 20, further evidence that condition *G^C^*5*G^T^* and the associated 371 genes may exhibit distinctive behaviour.

**Table 5:**
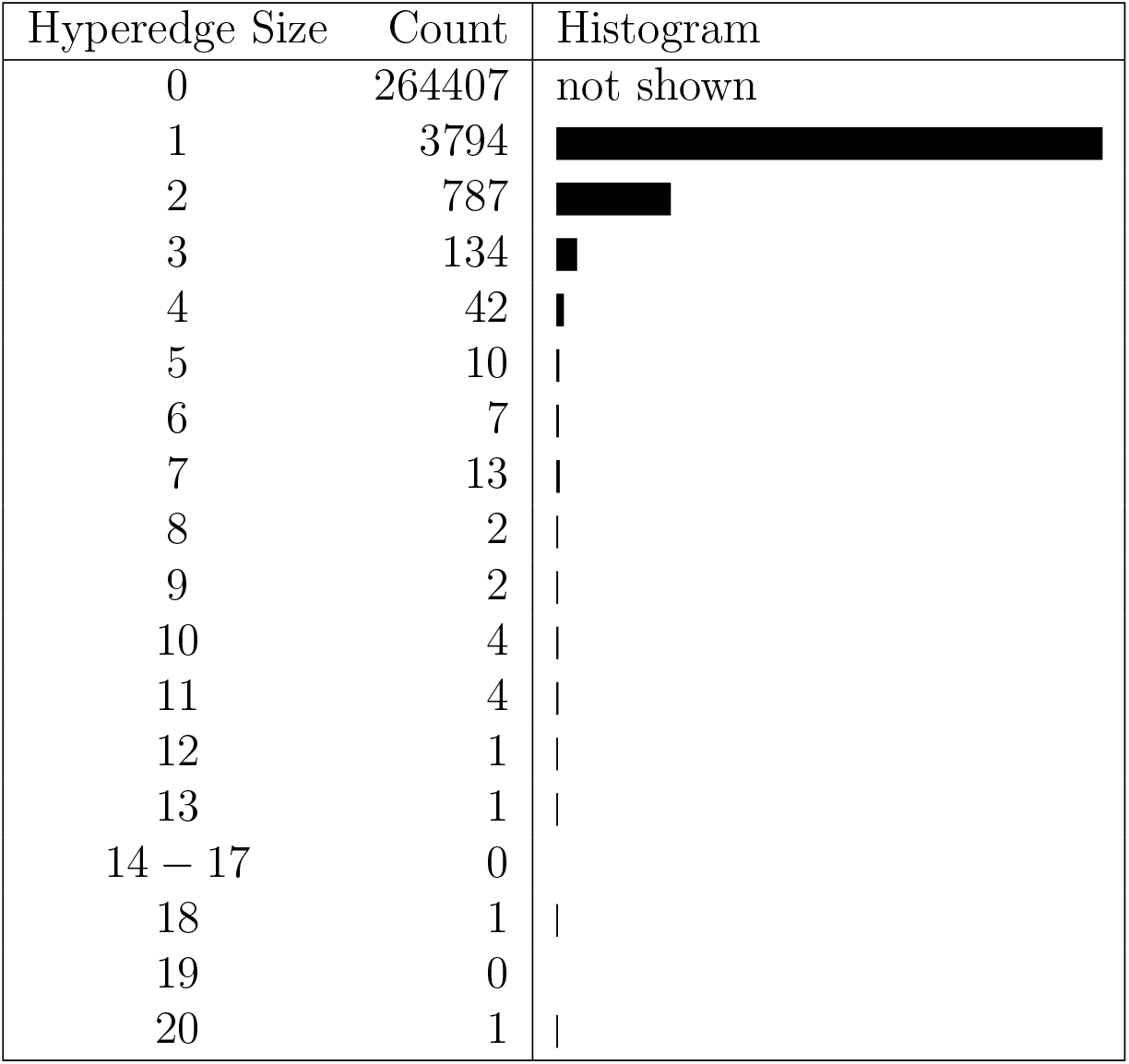
Hyperedge size distribution for Hypergraph 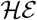.

| Hyperedge Size | Count | Histogram |
| --- | --- | --- |
| 0 | 264407 | not shown |
| 1 | 3794 |  |
| 2 | 787 |  |
| 3 | 134 |  |
| 4 | 42 |  |
| 5 | 10 |  |
| 6 | 7 |  |
| 7 | 13 |  |
| 8 | 2 |  |
| 9 | 2 |  |
| 10 | 4 |  |
| 11 | 4 |  |
| 12 | 1 |  |
| 13 | 1 |  |
| 14 – 17 | 0 |  |
| 18 | 1 |  |
| 19 | 0 |  |
| 20 | 1 |  |

While the above statistics provide good contextual information, this information is generally available without the need to define hypergraphs. The value of the hypergraph setup lies in highlighting the relationships between the hyperedges. That is, identifying sets of hyperedges that coincide with the same set of conditions, revealing patterns of behaviour within the gene set. Moreover, isolated hyperedges of size one, indicate genes that satisfy a single condition, a feature distinguishing them from the majority.

### 4.2. Line Graph Analysis

In this subsection we provide a more comprehensive study by focusing on line graphs for small values of *s*. We simplify the presentation using the *collapsed s*-line graph, in which sets of equivalent hyperedges are consolidated and represented with a single vertex. Thus each vertex represents an *x*-multiplicity hyperedge, and we say that the vertex has multiplicity *x*. Each vertex is labelled by its multiplicity, and the radius of the corresponding circle indicates the number of conditions that each hyperedge satisfies. One immediate advantage is to show the natural partitioning of the data into sets of equivalent hyperedges.

To illustrate this point, consider the 1-line graph for 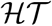 (Figure 2). It contains a single isolated vertex (hyperedge), a clique of order 11 and a densely connected component on 1133 vertices (hyperedges). Cross referencing with Table 2 identifies the isolated vertex as a single gene satisfying condition 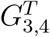 and the clique corresponding to a set of 11 genes satisfying only condition 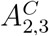. Given their distinct characteristics these 12 genes warrant further biological investigation, with results briefly discussed in Section 5.

**Figure 2:**
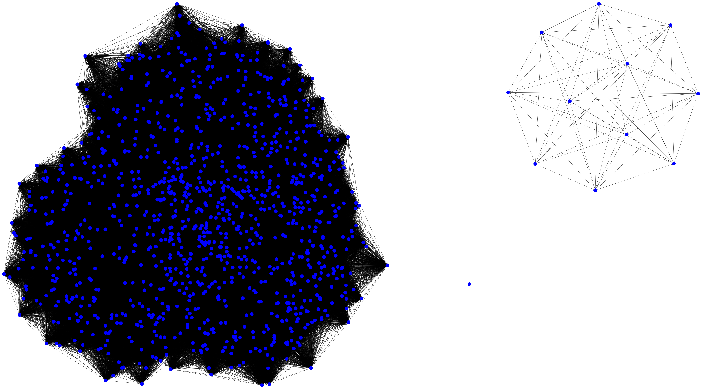
The 1-line graph representation of 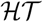, where vertices correspond to hyperedges (genes) and edges indicate at least one vertex (condition) in common.

As expected, the current visualisation does little to penetrate the large and dense component. This component is not a clique and has diameter two, thus there exists at least one pair of hyperedges not satisfying a common condition, but each sharing a common condition with a neighbour. Given that 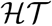 has many more hyperedges (genes) than conditions, it is reasonable to conjecture that 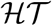 contains large numbers of hyperedges with large multiplicity, prompting an investigation of the collapsed 1-line graph, which is illustrated in Figure 3. The original 1133 vertices are consolidated into 18, with multiplicity between 2 and 219.

**Figure 3:**
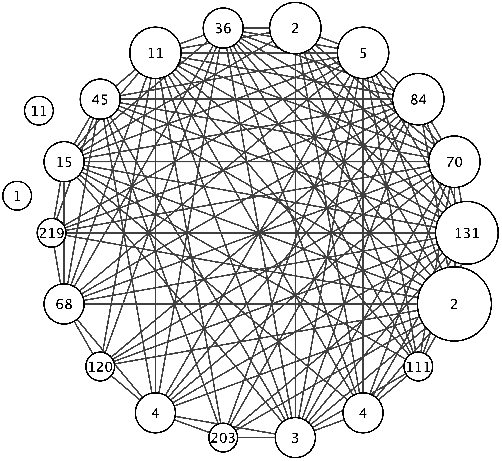
The collapsed 1-line graph representation of 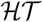, where a vertex is labelled with the multiplicity of the hyperedge (genes) and size of a vertex represents the size of the hyperedges (number of conditions).

Taking *s* = 2 suppresses the large number of hyperedges of size one in 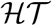 (Table 4) and we see that the collapsed 2-line graph (Figure 4) is connected and partitions the 480 hyperedges (size ≽ 2) into 14 collapsed vertices. This 2-line graph highlights a 2-multiplicity hyperedge (of size five incident with all high degree conditions 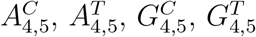, see Table 4), that forms a cut-vertex (shaded). This 2-multiplicity hyperedge shares at least two conditions with all other hyperedges, while sharing at least two other common conditions with a group of six hyperedges. The biological role of these genes will be discussed further in Section 5.

**Figure 4:**
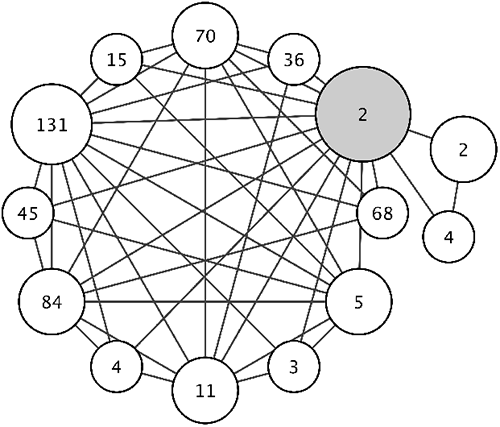
The collapsed 2-line graph representation of 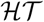, where a vertex is labelled with the multiplicity of the hyperedge (genes) and size of a vertex represents the size of the hyperedges (number of conditions). The cut-vertex is shaded.

Removal of the cut-vertex (Figure 4) reveals a large component of diameter two, that can be characterised as a set of hyperedges that contain various combinations of the high degree conditions 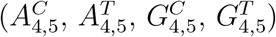, each with somewhat different probability, but little other discernible structure.

In 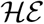, the number of non-empty hyperedges and correspondingly the number of vertices in the 1-line graph is 4803. These vertices are partitioned into 189 clusters (*x*-multiplicity hyperedges) in a densely connected collapsed 1-line graph. However, it has diameter three indicating that there exists at least one pair of hyperedges in 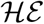 which do not satisfy common conditions, but in addition neither do their neighbours, so there is no linking hyperedge which intersects both. In terms of gene expression, this may suggests greater variation between genes.

The (collapsed) 3-line graph of 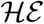, comprised of one major component and four isolated vertices, can be analysed as above. However, more interesting is an analysis of the hyperedges incident with condition *G^C^*5*G^T^*, an outlier identified in the degree sequence analysis. The four isolated vertices correspond to 4 distinct hyperedges, representing 34 genes in total, that satisfy condition *G^C^*5*G^T^*, a condition common to each of these 34 hyperedges but not incident with the single hyperedge that satisfies all 20 other non-zero degree conditions. The four isolated vertices are distinguished by the remaining incident conditions. That is, a 29-multiplicity hyperedge that also satisfies conditions *A^C^*5*G^C^*, *A^T^*5*G^C^*, a 3-multiplicity hyperedge that also satisfies conditions *A^T^*5*G^T^*, *A^C^*5*G^T^*, then two 1-multiplicity hyperedges that also satisfy, respectively, *A^T^*1*G^C^*, *A^C^*5*G^T^* and *A^T^*2*G^C^*, *A^T^*4*G^C^*. This latter hyperedge arises again in the subsequent discussion on the betweenness where it is identified as an outlier and distinguishing the associated subset of genes.

Notably in the major component there is a pendant vertex, of multiplicity 2, adjacent to a cut-vertex also of multiplicity 2, which in turn is adjacent to a further six 1-multiplicity vertices. Again we see that the condition *G^C^*5*G^T^* is prominent, occurring in the intersection of the pendent and cut vertices together with conditions *A^C^*5*G^T^*, *A^T^*4*G^C^*, but not in the pairwise intersection of the cut-vertex with the other six vertices, which is characterised by inclusion of conditions *A^C^*5*G^T^*, *A^T^*4*G^C^*, *A^C^*1*G^T^*. This suggests that although this set of eight hyperedges all share two conditions, it is the cut-vertex hyperedge pair which act as a linking hyperedge through particular conditions.

Investigating further the set of line graphs, we note that the major component of the 3-line graph has diameter three, or equivalently there exists a set of four hyperedges that form a 3-path. Since the hyperedge of size 20 is incident with all conditions except *G^C^*5*G^T^* it will intersect all hyperedges of size three or more except those of size three that satisfy condition *G^C^*5*G^T^*. Thus it is condition *G^C^*5*G^T^* that determines the structure of the paths of length three on the 3-line graph. This is further exemplified as the diameter of the major component is reduced to two when the pendent vertex is removed. The 4-line graph is comprised of two components, one consisting of a 2-multiplicity hyperedge (which again must be incident on *G^C^*5*G^T^*) with the remainder of the hyperedges being incident on a large dense component.

More interesting is the exception to the decline in distribution of hyperedge sizes in 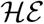 (Table 5), where the number of hyperedges of size seven increases to 13, suggesting the 7-line graph (Figure 7) may reveal more information. There are 29 hyperedges that satisfy seven or more conditions and pairwise most of these have distinct intersection patterns, with the exception of a 2-multiplicity hyperedge and a 3-multiplicity hyperedge. The conditions contained within the latter follow a distinct pattern, satisfying the second and fifth time point comparison for all agravitropic and gravitropic contrasts, with the exception of condition *A^C^*2*G^C^*. Thus, indicating that these three genes (hyperedges) reveal distinct and rare differential behaviour across phenotypes at particular time points. We note that this behaviour is only seen in the hyperedges of size 18 and 20.

**Figure 5:**
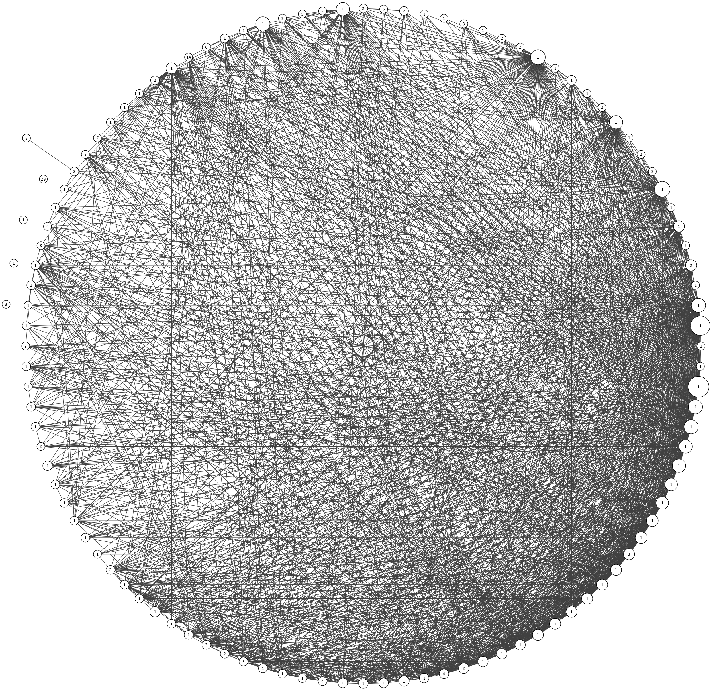
The collapsed 3-line graph representation of 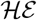, where a vertex is labelled with the multiplicity of the hyperedge (genes) and size of a vertex represents the size of the hyperedges (number of conditions).

**Figure 6:**
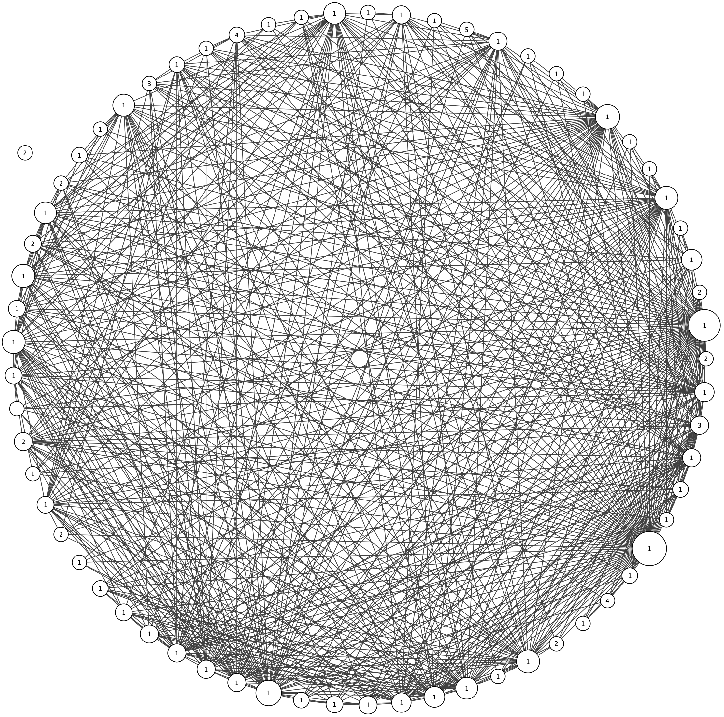
The collapsed 4-line graph representation of 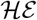, where a vertex is labelled with the multiplicity of the hyperedge (genes) and size of a vertex represents the size of the hyperedges (number of conditions).

**Figure 7:**
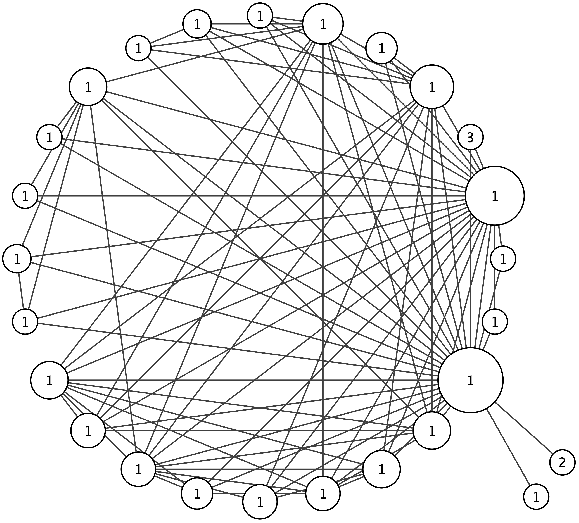
The collapsed 7-line graph representation of 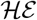, where a vertex is labelled with the multiplicity of the hyperedge (genes) and size of a vertex represents the size of the hyperedges (number of conditions).

The former, the 2-multiplicity hyperedge, with distinct intersection pattern, forms a pendant vertex and strictly center around comparisons between the agravitropic and gravitropic phenotype. However, unlike the collection of three hyperedges, the conditions satisfied have less emphasis on time steps. Specifically these hyperedges satisfy conditions *A^C^*2*G^T^*, *A^T^*2*G^C^*, *A^T^*2*G^T^*, *A^C^*3*G^T^*, *A^T^*3*G^T^*, *A^C^*5*G^C^*, *A^C^*5*G^T^*.

In the next section we extend the current approach and analyse centrality measures on the line graphs to rank the hyperedges in the large dense components.

### 4.3. Centrality Analysis

While the collapsed line graph can reduce some of the complexity for hypergraphs of high order and size (for example 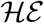) some structural features of the data remain opaque. Thus we further explore centrality measures such as closeness, betweenness and eigencentrality as tools for identifying subsets of hyperedges that exhibit similar behaviour, as well as outliers (see for instance Figures 10, 11 and 12). To emphasise the validity of this analysis, we begin by studying the collapsed 2-line graph (Figure 4) for 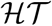, verifying that the chosen centrality measures can identify salient features as documented above. Once this is established we will use centrality measures to reveal more of the structure of 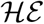.

Recall that for graphs of diameter less than three the closeness tracks the degree, so in Figures 8 and 9, we tabulate only the betweenness and eigencentrality for the collapsed 2-line graph of 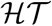.

**Figure 8:**
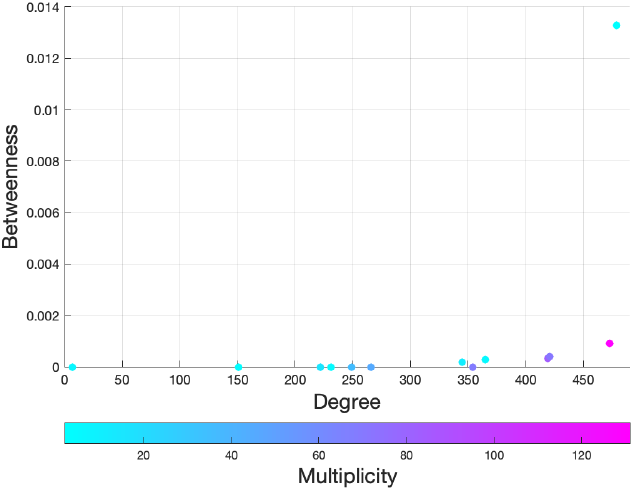
The betweenness and degree scores of the 2-line graph of 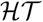.

**Figure 9:**
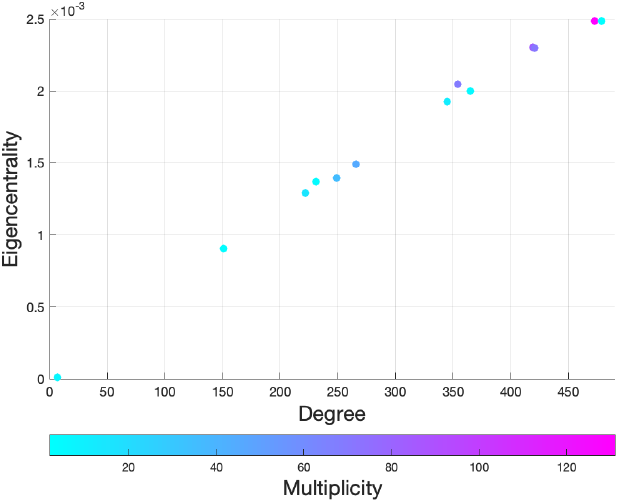
The eigencentrality and degree scores of the 2-line graph of 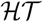.

In both plots, the outlier with high degree, betweenness and eigencentrality, is the cut-vertex which was identified as a 2-multiplicity hyperedge in the collapsed 2-line graph. Since the diameter of the 2-line graph is two, it is not surprising the betweenness centrality will distinguish this 2-multiplicity hyperedge (genes) of size five that share common characteristics with the major and minor subsets of hyperedges (genes). In contrast, the eigencentrality is high for both the cut-vertex (see Figure 4) and the next highest degree vertex, corresponding to a 131-multiplicity hyperedge, all incident with conditions 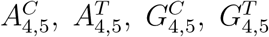. The distinction is that the cut-vertex also satisfies condition 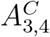. There are only six other hyperedges (genes) which satisfy this condition, stratified into a 4-multiplicity hyperedge and 2-multiplicity hyperedge. From this, we deduce that condition 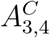 does not significantly contribute to the eigencentrality of the cut-vertex. This is further exemplified, as the removal of the cut-vertex marginalises six hyperedges of low degree and as expected, vertices in this minor component have lower relative eigencentrality (see Figure 9).

Having established that these centrality measures can identify notable structure in hypergraphs, we turn our attention to 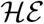, in which direct analysis of the line graphs gave limited insight. We begin by studying the collapsed 1-line graph, which is of diameter three. As noted earlier the closeness will track the degree distribution in the main. However, Figure 10 does indicate the existence of a subset of lower degree vertices which deviate from the trend. Specifically, identifying vertices with eccentricity three (distance three from at least one other vertex) and in terms of genes, identifying a subset of genes that exhibit atypical characteristics, in that they have distinct profiles with respect to the contrasts.

**Figure 10:**
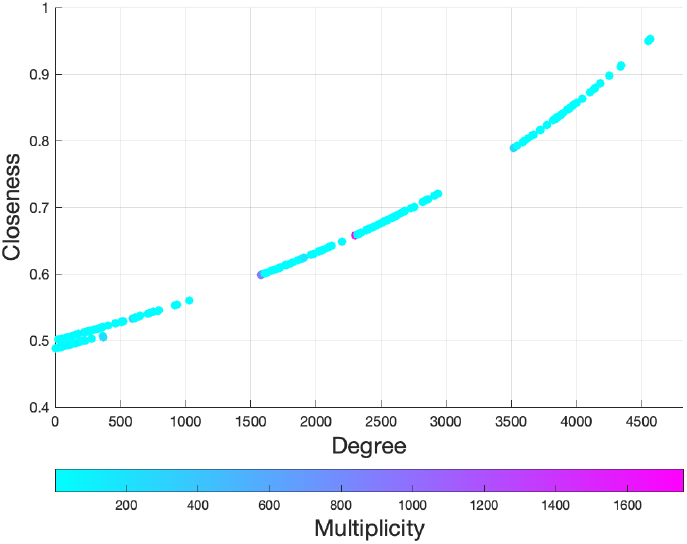
The closeness and degree scores of the 1-line graph of 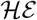.

In combination, the betweenness and eigencentrality provide a rich source of structural information about 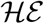; see Figures 11 and 12. The vertices are partitioned into three groups by degree, but in combination with eigencentrality we see an improved partition into four groups (*P*_1_, *P*_2_, *P*_3_, *P*_4_). Betweenness appears to provide complementary information, distinguishing vertices within these clusters.

**Figure 11:**
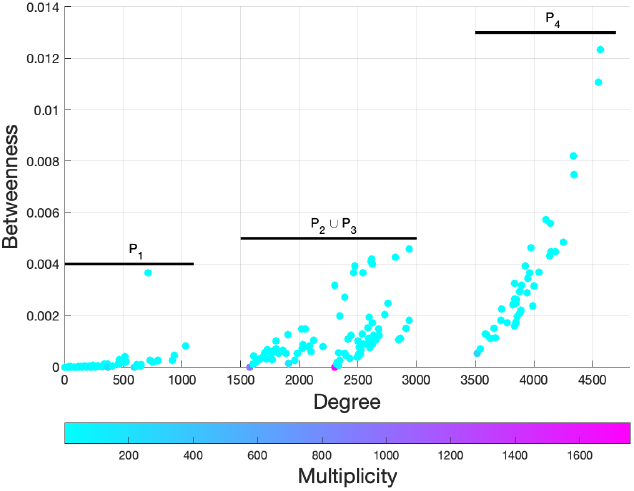
The betweenness and degree scores of the 1-line graph of 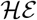.

**Figure 12:**
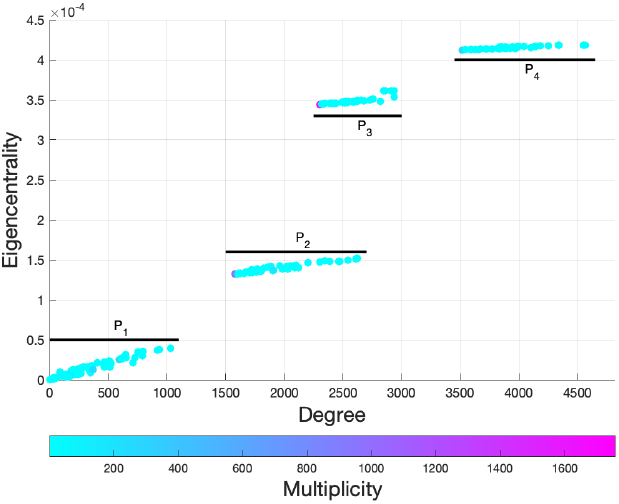
The eigencentrality and degree scores of the 1-line graph of 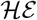.

Both measures show insignificant centrality scores for vertices (hyperedges) with degree in partitions *P*_1_, except for one outlier. This outlier, with an increased betweenness score, corresponds to a 1-multiplicity hyperedge satisfying three conditions *G^C^*5*G^T^*, *A^T^*2*G^C^*, *A^T^*4*G^C^* and intersecting 76 subsets of collapsed hyperedges (712 non-collapsed hyperedges). Note the inclusion of the condition *G*^C^5*G^T^*, which was discussed earlier as being a condition of interest. Roughly, this betweenness result indicates that the fraction of shortest paths incident with this vertex, is at least double the corresponding fraction for all vertices in the same degree partition. We note that this betweenness is among the largest across all vertices and is not distinguished by the eigencentrality. Indeed there is a large collection (denoted here by *α*) of vertices, each of degree 1579 and betweenness zero.

Investigating further we have identified a cluster, being a 910-multiplicity hyperedge that satisfies the single condition *A^C^*5*G^C^*. The degree sequence in Table 3 implies the existence of another 670 hyperedges (partitioning into 79 subsets) satisfying this condition. Within the collection of 1580 hyperedges, the 910-multiplicity hyperedge in question has the smallest degree. Thus these vertices will not be incident with any shortest path in the 1-line graph, justifying a betweenness score of zero. An identical situation can be found for a collection (denoted here by *β*) of vertices of degree 2305, which only satisfy condition *A^C^*5*G^T^*.

For vertices in the partition *P*_2_ ∪ *P*_3_, there is an increased variation in betweenness scores compared to those in partition *P*_1_. This variation is carried into the eigencentrality scores, which partitions this collection of vertices into distinct groups (*P*_2_ and *P*_3_). It is interesting to note that the lowest degree vertices in each of the eigencentrality partitions (*P*_2_ and *P*_3_), are contained within the collections *α* and *β*. So the eigencentrality creates partitions based on the clusters, emphasising different clusters with respect to the minimum degree vertices (*α* and *β*).

Vertices with the same degree (overlap in *P*_2_ and *P*_3_) have distinctly different eigencentrality scores. Given that the eigencentrality reflects the number of times a vertex is traversed in an infinite random walk on the graph, this indicates groups of hyperedges (vertices) with similar intersection patterns but different roles within the 1-line graph. In addition, this information is not necessarily reflected in the betweenness scores. In the final partition for both measures (*P*_4_), we may use betweenness to identify extreme cases where vertices attain extreme scores against the trend. Figure 11 highlights two outlier vertices, being the hyperedges of greatest size, that is, sizes 18 and 20 (Table 5). Consequently, most hyperedges will intersect these two, explaining the large betweenness scores.

Delving further, we can see that the four partitions, *P*_1_, *P*_2_, *P*_3_, *P*_4_ can be differentiated by the inclusion of hyperedges satisfying none, one or both of the conditions *A^C^*5*G^C^* and *A^C^*5*G^T^* (Table 2). Specifically, hyperedges in *P*_1_ satisfy neither and have low betweenness and eigencentrality, hyperedges in *P*_4_ satisfy both and have high eigencentrality. The subset of hyperedges in *P*_2_ are characterised by hyperedges that satisfy condition *A^C^*5*G^C^* but not *A^C^*5*G^T^* with relatively low eigencentrality and the subset of hyperedges *P*_3_ satisfy *A^C^*5*G^T^* but not *A^C^*5*G^C^* with higher eigencentrality. This supports our initial supposition that these two conditions are central to the analysis and act as hubs of the hypergraph. This result suggests that in some cases analysis of the centrality of the vertices (hyperedges) in the line graph and the corresponding results can be linked not only to the hyperedges in the hypergraph but also to the centrality measures on its vertices.

We may deduce that the structure of 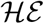 is dominated by the intersection of two large clusters of hyperedges or two high degree vertices in the collapsed 1-line graph. The larger cluster is defined by the inclusion of condition *A^C^*5*G^T^*, a condition that appears to be more central to the overall structure of 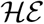, as indicated by the higher eigencentrality score for hyperedges in *P*_3_ than in *P*_2_, even when the latter have higher degree. The other cluster is characterised by the inclusion of *A^C^*5*G^C^*, but with hyperedges containing this condition being less central to the structure of 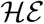. This is consistent with the setup of the experiment, where conditions were designed to test the differentiation in expression levels for the gravitropic response.

Given our understanding of the broad structure of the 1-line graph, betweenness analysis allows us to identify hyperedges which correspond to potentially significant genes. With this, we see that the centrality scores were able to reveal multiple salient features about the hypergraph. Degree alone gave a natural partitioning into 3 groups, but eigencentrality provided a much clearer delineation of the major structure, showing a partition into 4 distinct groups. Betweenness then allowed us to identify hyperedges of interest within each group, identifying potentially interesting genes and gene associations. While eigencentrality allowed for differentiation between same size hyperedges, distinguishing genes based on key conditions.

## 5. Biological Significance of Findings

The analysis and corresponding results identified within the hypergraph framework have illuminated significant features within the data, identifying a variety of genes (candidate gene sequences) that show functional variation across differing aspects of the study. In this section, we explore the biological significance of these genes and discuss possible further investigations.

Significant genes identified through hypergraph 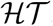, and the associated time related conditions (Equation 4) include two outlier genes. These genes have the largest size, each satisfying five conditions (Table 4), and act as cut-vertices in the the 2-line graph (Figure 4), with both facts leading to a significantly higher betweenness score (Figure 8). In particular, as cut-vertices they act as a natural link which when removed partition, into two subsets, the set of all genes satisfying at least two conditions. These facts emphasise the distinctive nature and significance of these genes, where investigations reveal these genes as predictive indicators. Further, both genes were annotated as REVEILLE1 (RVE1)-like genes, which is a type of Myb-like transcription factor that integrates the circadian clock and the auxin pathway [26]. RVE1 genes have also been found to have functions involved in the stimulation of plant growth by interacting with auxin biosynthesis genes and promoting cell elongation [26]. As the expression of these circadian rhythm genes was found to be significant in time-related conditions it may imply that these genes control differences both in gravitropic and circadian responses between the gravitropic and agravitropic populations.

The removal of the cut-vertices from the 2-line graph identifies a small subset of six genes. These six genes and the two cut-vertex genes exhibit unique expression profiles in that they are the only genes to satisfy condition 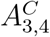. These six genes were annotated within the ARABIDOPSIS PSEUDORESPONSE REGULATORS (APRR) family of genes. APRR genes are known to be involved in circadian rhythm and elongation during seedling growth [27].

It is also worth remarking that the 1-line graph of 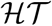 (Figures 2 and 3) highlights a clique of 11 genes, satisfying only 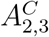. The majority of these genes were found to be chlorophyll a-b binding, which is involved in photosynthetic regulation [28]. This 1-line graph also identifies an anomaly, a single gene with a unique expression profile (satisfying 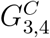) showing no commonality with any other gene exhibiting significant differential expression. It is difficult to interpret this singularity in the gravitropic control group, but we remark that this gene was able to be annotated against the *Arabidopsis* protein database (Tair Blast) and was found to be involved in the biosynthesis of isoleucine. Isoleucine is a vital amino acid as it is involved in energy production, and has also been shown to be involved in abiotic stress responses [29].

We now move onto hypergraph 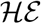, where two outlier genes are prominent in the hyperedge size analysis (Table 5), the betweenness (Figure 11) and the eigencentrality (Figure 12) of the 1-line graph. Individually these genes satisfy 20 and 18 conditions and, respectively, were identified as a ubiquitin gene and a polyubiquitin gene. The exact annotation of these genes is unclear, however the ubiquitin-like gene could be annotated against the Arabidopsis transcriptome as RELATED TO UBIQUITIN 1(RUB1) gene, also known as *NEDD8* [30]. Mergner and Schwechheimer [31] identified components of the neddylation and deneddylation pathways based on mutants with defects in auxin and light responses. The characterization of these mutants was instrumental for the elucidation of the neddylation pathway, which appears to function as a stable post-translational protein modifier. An AMP-RUB1 intermediate is formed by an activating enzyme, distinct from the ubiquitin activating enzyme E1, which is composed of a heterodimer AXR1/ECR1 [32]. Auxin response is mediated, at least in part, through modification of cullin AtCUL1 by the attachment of RUB1 to ‘Lys-692’ [33]. These results suggest the possible involvement of calmodulin in auxin-induced elongation. RUB1 genes have been shown to play a vital role in Arabidopsis, with roles relating to vegetative growth, auxin signalling and ethylene production [34]. The identification of this gene could provide a candidate for further investigation into its role during the evolution of gravitropic differentiation in *S. lautus*.

The next gene we explore is the outlier which can be identified in Figure 11, specifically in partition *P*_1_ for the 1-line graph of 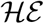. This gene was found to have a betweenness score over double all other genes with similar degree scores. This gene is not differentiated in the eigencentrality nor in the visual representation of the dense and complex 1-line graph. This gene was annotated to the *A. thaliana* SHAGGY-LIKE KINASE GROUP 2 (SK2-2) also known as BRASSINOSTEROID INSENSITIVE 2-like, which is involved in the brassinosteroid-mediated signalling pathways [35], related to light-regulated hypocotyl elongation [36] and multiple stress responses including responses to drought [35]. SK2-2 has eight gene interactions of which most are calmodulin genes. Yang and Poovaiah [37] provided the first direct evidence for the involvement of calcium/CaM-mediated signaling in auxin-mediated signal transduction and [38] where Poovaiah Yang and van Loon established that Calcium and calmodulin (CaM) play an important role in gravity signal transduction mediating gravitropic response in plants.

The final collection of genes which we will discuss is the clique of 29 genes identified in the 3-line graph of 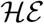, as per Figure 5. This is the largest cluster of genes found in this line graph, with the next largest being only 14 genes. We conclude that the expression profiles for these 29 genes are similar, while distinctly different from all other genes that satisfy at least three conditions. Investigations show that tubulin genes are well-represented in this group (15 out of 29 genes). Tubulin is a protein that contributes to the formation of microtubules, involved in cell division, polarity of growth, cellwall deposition, intracellular trafficking and communications [39]. They also allow for plants to increase the rigidity of their cell walls [40]. Additionally, [41] and [42] found that cytoskeletal elements (such as actin microfilaments and microtubules) are important for pressure perceiving systems, such as sensing gravity. The importance of plant cytoskeletons in the early stages of gravity response signalling are given by Kiss in [43], subject to the caveat that the exact mechanisms of this process remain elusive.

This analysis has uncovered genes relevant to the interpretation of gravitropic responses in *S. lautus* genes involved in circadian rhtyhm responses, which are expected to be observed in time-course experiments. Through both hypergraph frameworks, we were able to identify genes and related functions significant in the phenotypic gravitopism responses. We also uncovered some genes exhibiting unique expression profiles with the known function suggesting further investigation is warranted.

## 6. Conclusion

In this paper, we have documented research that validates hypergraphs as a powerful framework for the study of biological data sets. While our work has foundation that is presented in [5] and other studies, our methods differ in multiple ways. Firstly, the hypergraph construction presented here combines fold change and *p*-values, reducing the family-wide false discovery rate, creating a more robust hypergraph, leading to more meaningful results. This strict selection criteria for the inclusion of a condition within a hyperedge reduces noise and complexity, thus enhancing access to a range of analytical techniques. Secondly, further reductions in complexity were achieved by resolving the list of test conditions into two groups, giving a hypergraph focused on time comparisons and another hypergraph based on experimental group comparisons. Lastly, we analysed hypergraphs using both analytical and visual techniques. We first looked at the degree sequence and size distribution to gauge the structure of the hypergraph. This shed light on trends and outliers in the data which lead to biological investigation.

Robust distance measures were obtained by representing hypergraphs in terms of **s**-line graphs. The collapsed version was particularly useful as it was able to identify clusters and cliques of genes with similar and unique behaviour and as such were useful visual tools. This definition of distance enabled the calculation of multiple centrality measures, with particular emphasis on betweenness and eigencentrality. The information gained here was not readily available from an analysis of the degree sequence or the hyperedge size, and provides additional validation for the introduction of the line graphs, the subsequent distance measures and ultimately the analysis of the centrality measures.

Overall, these analysis techniques did highlight key information about genes which exhibited distinct expression profiles, even in dense and complex data. In summary, the hypergraphs and associated techniques discussed in this paper were found to be useful in highlighting salient features within the data set.

